# Integrin-mediated attachment of the blastoderm to the vitelline envelope impacts gastrulation of insects

**DOI:** 10.1101/421701

**Authors:** Stefan Münster, Akanksha Jain, Alexander Mietke, Anastasios Pavlopoulos, Stephan W. Grill, Pavel Tomancak

## Abstract

During gastrulation, physical forces reshape the simple embryonic tissue to form a complex body plan of multicellular organisms^1^. These forces often cause large-scale asymmetric movements of the embryonic tissue^2,3^. In many embryos, the tissue undergoing gastrulation movements is surrounded by a rigid protective shell^4,5^. While it is well recognized that gastrulation movements depend on forces generated by tissue-intrinsic contractility^6,7^, it is not known if interactions between the tissue and the protective shell provide additional forces that impact gastrulation. Here we show that a particular part of the blastoderm tissue of the red flour beetle *Tribolium castaneum* tightly adheres in a temporally coordinated manner to the vitelline envelope surrounding the embryo. This attachment generates an additional force that counteracts the tissue-intrinsic contractile forces to create asymmetric tissue movements. Furthermore, this localized attachment is mediated by a specific integrin, and its knock-down leads to a gastrulation phenotype consistent with complete loss of attachment. Moreover, analysis of another integrin in the fruit fly *Drosophila melanogaster* suggests that gastrulation in this organism also relies on adhesion between the blastoderm and the vitelline. Together, our findings reveal a conserved mechanism whereby the spatiotemporal pattern of tissue adhesion to the vitelline envelope provides controllable counter-forces that shape gastrulation movements in insects.

---

It is not birth, marriage or death but gastrulation that marks the most important event in everyone’s life, as Lewis Wolpert famously quipped. Animal gastrulation is characterized by the transformation of a single-layered blastula into a multi-layered gastrula consisting of so-called germ-layers from which all future embryonic tissues develop. This transformation is accompanied by large-scale tissue flows and folding events that are thought to be driven largely by tissue-intrinsic contractile forces^8–17^. In most species, gastrulation occurs within some form of a rigid shell surrounding the developing embryo, yet the interaction of the living tissue with the inner surface of the surrounding shell and the role of this interaction for gastrulation has not been explicitly considered.

In the red flour beetle *Tribolium castaneum*, the cellular blastoderm is confined by the vitelline envelope and remains closely apposed to it while undergoing large morphogenetic movement during gastrulation^18,19^. This makes it an exquisite model for studying how gastrulation is impacted by interaction with its surroundings. In this system, gastrulation begins when approximately 2/3 of the blastoderm folds inwards to form the embryo that gets entirely engulfed by the remaining 1/3 destined to be become the extra-embryonic serosa (**Fig 1a**). This morphogenetic event is characterized by large-scale posterior flow of the dorsal tissue, indicating that the blastoderm is rather free to slide underneath the vitelline envelope. At the same time, the tissue at the anterior-ventral side remains stationary. Hence, the blastoderm exhibits large-scale unidirectional flow (**Fig. 1a**). However, such asymmetric movement is difficult to reconcile with current models of tissue mechanics where movements are thought to arise exclusively through contractile force generation within the tissue^14,20–22^. This is because inertial effects are negligible in biological systems and therefore tissues will only move when forces are acting upon them. Moreover, in a closed system consisting of a tissue that does not interact with its environment forces add up to zero, and one would expect that the tissue flows driven by these forces also add up to zero (see SI). Hence, tissue-intrinsic contractile forces alone should not be able to generate unidirectional flow of the Tribolium blastoderm.

**Figure 1:**
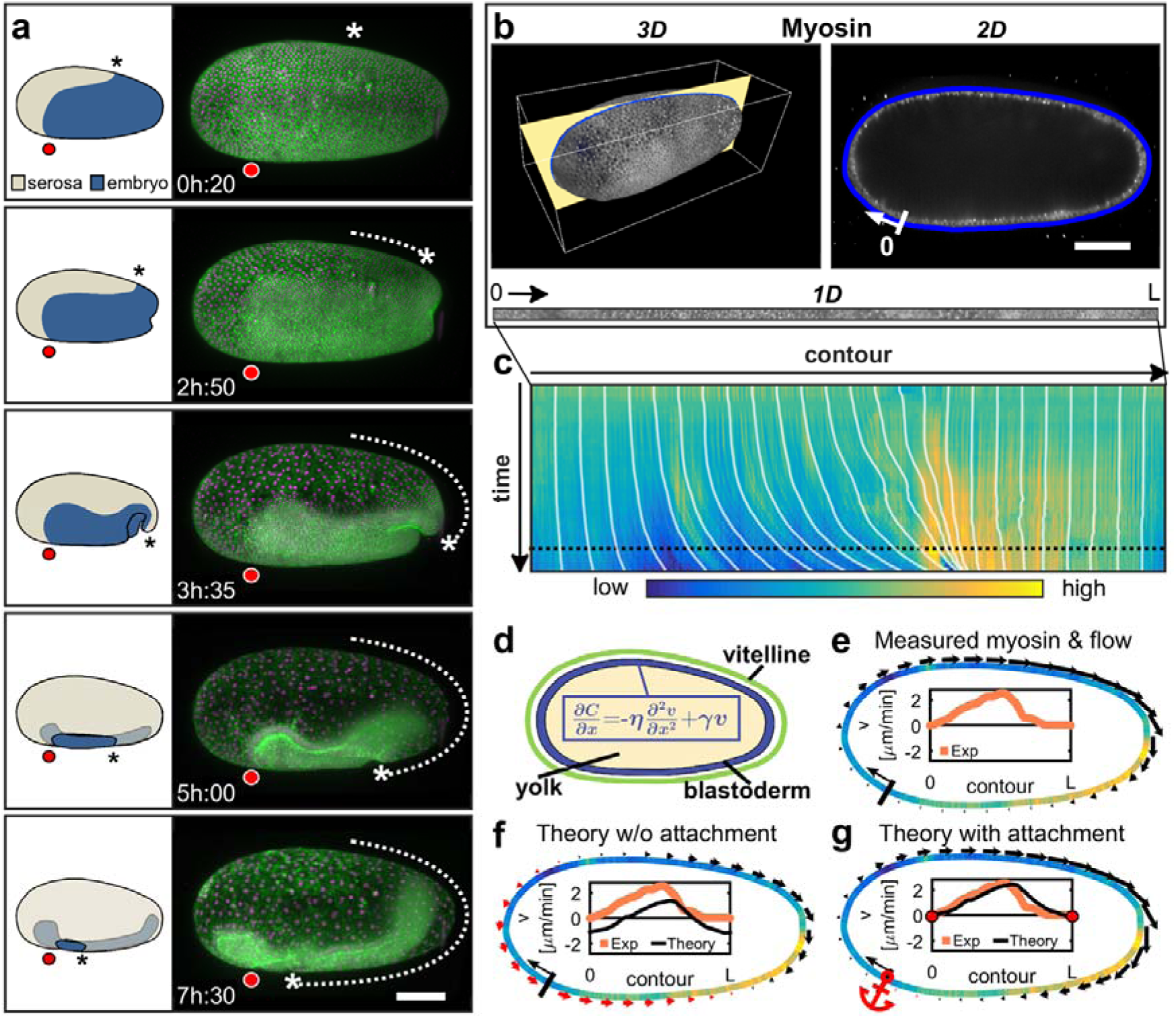
Unidirectional tissue flow during early gastrulation of *Tribolium castaneum.* (**a**) Lateral view of a gastrulating Tribolium embryo in a schematic (left) and in a maximum intensity projection of a line transiently expressing LifeAct-eGFP (green) and Histone2A-mCherry (magenta) imaged by time-lapse light sheet microscopy (right). The serosa moves unidirectionally towards and across the posterior pole (dashed arrow and asterix) while the anterio-ventral part remains stationary (red circle). Scale bar: 100 μm. (**b**) Right panel: 2D sagittal cross-section taken from 3D reconstructed multi-view light sheet recording (yellow plane in left panel) of embryo transiently expressing Tc.sqh-eGFP. Blue outlines indicate the vitelline envelope. Scale bar in 2D panel: 100 μm. Bottom panel: unwrapped 1D contour opened at the origin 0. (**c**) Kymograph of the evolution of the myosin intensity along the 1D contour in **b** over the course of gastrulation. White lines show flow measured by PIV tracking. The horizontal dashed line denotes the time point displayed in **e-f**. (**d**) Schematic representation of a cross section through a Tribolium embryo. Inset: equation of motion of the tissue using thin-film active gel theory. (**e-f**) Experimentally determined myosin intensity distribution along the the sagittal contour (color map as in **c**), and measured (**e**) or predicted (**f,g**) tissue flow velocity fields at a single time point (black arrows denote positive and red denote negative flow velocities). Insets: flow velocity plotted over embryo circumference. (**e**) Experimentally determined flowfield. (**f**) Theoretically predicted flowfield, assuming that the tissue flows freely with respect to the vitelline envelope. (**g**) Theoretically predicted flowfield, assuming the tissue is anchored at the anterio-ventral side of the embryo (red anchor). Note that in contrast to (**f**), the predicted flowfield matches the measured one (compare insets).

To resolve this apparent paradox, we sought to investigate the forces governing unidirectional morphogenetic tissue flow in Tribolium. In many organisms, non-muscle myosin II (hereafter referred to as myosin) activity is the dominant mechanism of contractile force generation in epithelia ^14,15,17,21,22^. Thus, we first set out to evaluate the contribution of myosin-dependent contractile forces to the generation of unidirectional flow. To this end, we quantified the tissue flow and the myosin distribution by *in toto* multi-view light sheet imaging^23–26^ of the regulatory light chain of myosin, visualized through transient expression of Tc.sqh-eGFP (*spaghetti squash*, TC030667). Despite the complexity of this morphogenetic process^18^, key features of Tribolium tissue flows are captured in the mid-sagittal cross-section of the embryo where the blastoderm flows exclusively parallel to the circumference (**Fig. 1b** & Supplementary Video 1). Along this 1D contour, we obtained the distribution of myosin intensity over time and extracted the tissue flow field via particle image velocimetry (**Fig. 1c,e**). Because the blastoderm forms a relatively thin layer compared to the dimensions of the egg (**Fig. 1d**), we made use of thin-film active gel theory^27–29^ to predict the tissue flow-field generated by the spatial variation of myosin activity in the tissue^14,20–22^. This theory relates tissue flow to the global myosin distribution through two material parameters that determine how far active mechanical forces propagate inside the tissue^30^: friction between the tissue and its surroundings, and internal viscosity of the tissue layer. We determined the parameters for which the flow-field calculated from the theory best matches the experimentally measured one (see SI). While we find good agreement between the flow-fields from theory and experiment on the dorsal side of the embryo, the theory predicts the ventral tissue to flow posteriorly, a feature never observed in experiments (**Fig. 1f** & Supplementary Video 2). The theoretical prediction can be intuitively understood by examining the distribution of myosin along the embryo circumference: initially, myosin is distributed uniformly all over the embryo and no tissue flow is expected. However, at the onset of gastrulation, myosin becomes enriched in the embryonic part of the blastoderm (the posterior-ventral region) and concurrently becomes reduced in the serosa (the anterior-dorsal region) (**Fig. 1c,e** & Supplementary Videos 1,2). This global difference in myosin is expected to create flows towards the area of myosin enrichment from both sides: posterior flow on the dorsal side as well as posterior flow on the ventral side. Such symmetric flows are a consequence of the argument above that the sum of all flows, driven by forces that add up to zero, is also zero. Hence, myosin mediated tissue-intrinsic forces alone do not explain the stationary position of the anterior-ventral blastoderm portion. Therefore, additional forces must contribute to the observed tissue flows.

The absence of anterior ventral flow provides an important hint towards the source of such a force. For instance, it is conceivable that the anterior-ventral blastoderm interacts mechanically with the vitelline envelope^18,19,31^ ‘anchoring’ this region of the blastoderm in place. Such a local interaction would provide an additional external force that counteracts the forces generated by the tissue-intrinsic myosin activity. To investigate this hypothesis in our model, we included an anterior-ventral anchor point with infinite friction between tissue and vitelline envelope in the theory (see SI). This amended theory closely matches the experimentally measured magnitude of the large-scale dorsal tissue flow, while the ventral tissue remains stationary (**Fig. 1g** & Supplementary Video 3). Therefore, blastoderm anchoring to the vitelline in the anterior-ventral region together with myosin contractility of the tissue can quantitatively account for the observed unidirectional morphogenetic flow in Tribolium.

Next, we investigated this theory-based prediction, that the blastoderm is anchored to the vitelline, with several different experimental approaches. From earlier scanning electron microscopy studies it is known that microvilli protrude from the apical surface of the blastoderm all over the embryo^19^, and it is conceivable that these protrusions attach to the vitelline envelope^32^. We examined the distance between microvilli and the vitelline using transmission electron (TEM) tomography. Indeed, we observed that the ends of the protrusions are in direct contact with the vitelline envelope (**Fig. 2a,b**). Furthermore, we found small electron dense structures that could represent trans-membrane protein complexes at these connections (**Fig. 2b’**). We next imaged the dynamics of the apical protrusions via high-resolution live microscopy of Tribolium line constitutively expressing LifeAct-eGFP^33^ (**Fig. 2a,d** & Supplementary Video 4). We observed that the apical surface of the blastoderm cells is initially well separated from the vitelline envelope and the microvilli are bridging this space (**Fig. 2a,e**). This observation is consistent with the tissue being relatively free to move inside the vitelline at the onset of flows. However, preceding the onset of tissue flow, the protrusions of some cells on the ventral side of the embryo shrink and the apical surfaces of the cells get closer to the vitelline envelope, thus increasing their contact area (**Fig. 2a,c,e** & Supplementary Video 5). This raises the possibility that specific cells towards the anterior of the embryo may increase the strength with which they adhere to the vitelline.

**Figure 2:**
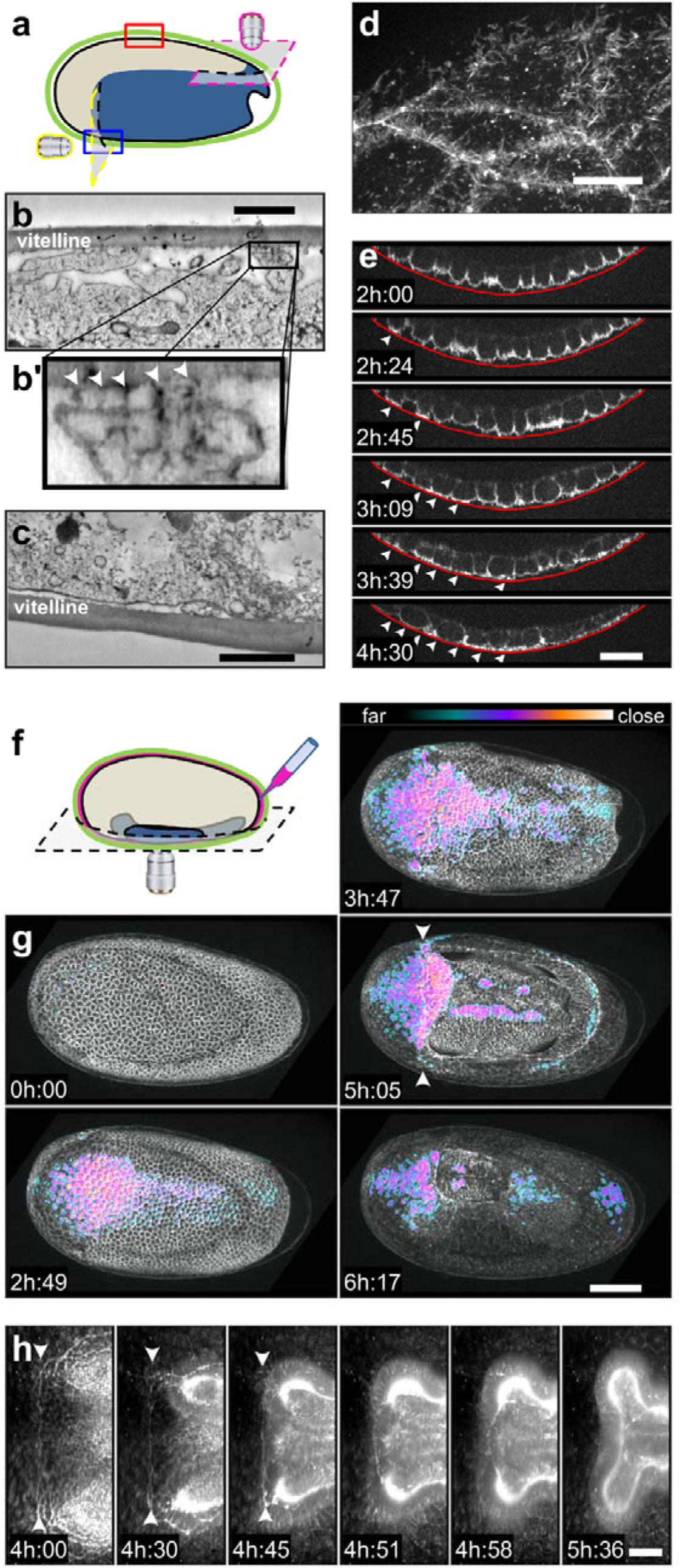
Attachment of the blastoderm to the vitelline envelope. (**a**) Schematic overview of the imaging fields of view and orientations around the Tribolium embryo shown in **b** (red box), **c** (blue box), **d** (magenta plane) and **e** (yellow plane). (**b**) Slice of a TEM tomogram depicting apical protrusions contacting the vitelline envelope. Scale bars in **b,c**: 500 nm. (**b’**) Magnified view of the region outlined in **b**. Arrowheads indicate electron dense material between the tip of the protrusion and the vitelline envelope. (**c**) Slice of a TEM tomogram depicting ventral side of the embryo. (**d**) Apical protrusions on the dorsal side of the embryo imaged by confocal microscopy during flow. Scale bar: 10μm. (**e**) Crosssection of a time-lapse recording acquired from the onset tissue flow at the anterior ventral side of a Tribolium embryo expressing LifeAct-eGFP. The red line indicates the position of the vitelline envelope. Scale bar: 20μm. (**f**) Schematic overview of the dextran (red) injection experiment. (**g**) Ventral view of an embryo expressing LifeAct-eGFP after rhodamine-dextran was injected into the perivitelline space. The dextran intensity signal was inverted and contrast-adjusted to yield a colormap representing the proximity of the apical blastoderm to the vitelline envelope (colorbar). Arrowheads mark the straight border between serosa and embryo. Scale bar: 100μm. (**h**) Ventral view of a map-projected light sheet recording at the late stage of gastrulation when the serosa window closes. Arrowheads mark the actin-rich straight border between serosa and embryo. Scale bar: 50μm.

To localize the exact group of cells that increase their proximity to the vitelline in space and time, we directly visualized the separation distance between the blastoderm and the vitelline. For this, we injected a solution of rhodamine-labelled dextran into the perivitelline space of a LifeAct-eGFP embryo (**Fig. 2f,g**). After the solution was injected at the posterior pole prior to the onset of gastrulation, the fluorescent signal distributed homogeneously across the perivitelline space surrounding the whole embryo (**Fig 2g** & Supplementary Video 6). This indicates a similar distance between blastoderm and vitelline all around the embryo. However, over the course of gastrulation, a restricted region at the anterior-ventral border between serosa and embryo exhibited a strong displacement of the dextran solution indicated by a local loss of intensity (**Fig. 2g** & Supplementary Video 6). Indeed, it is this very region of the blastoderm that remains stationary during gastrulation. Temporally, the formation of the contact zone defined by dextran exclusion precedes the onset of tissue flow and persists for ~4 hours indicating the formation of a stable attachment between the blastoderm and the vitelline. However, the dextran exclusion area eventually disappears at a later stage (**Fig. 2g**), and the dynamics of the tissue at these later time points provides additional evidence for anchoring of the blastoderm to the vitelline. During the final stages of serosa tissue closure around the internalized embryo, the border between embryo and serosa shows an upregulation of actomyosin resembling the supra-cellular actin cable^34^ seen during dorsal closure of Drosophila^35^. On the anterior-ventral side of the embryo, overlapping with the dextran exclusion zone, the cable is initially straight and stationary before it suddenly rips-off of at the edges and rounds up while the serosa window continues to close (**Fig. 2h** & Supplementary Video 7). When visualized as a sagittal cross-section it appears as if the blastoderm is forcibly peeled away from the vitelline envelope (see Supplementary Video 8). Taken together, these data indicate that the blastoderm attachment is localized to the anterior ventral region, that it is formed dynamically over time, and that high forces acting on this blastoderm region can eventually break its adhesion to the vitelline envelope.

We next tested directly whether this attachment is bearing a force during flow by disrupting it prematurely. If the attachment is mediated by protein complexes as suggested by the TEM data, digestion with a protease introduced into the perivitelline space should lead to an immediate disruption of adhesion. We would then expect the myosin-generated tissue contractility to pull the attachment zone posteriorly. To test this, we injected a solution of trypsin into the perivitelline space of an embryo during flow and observed the reaction of the ventral blastoderm (**Fig. 3a**). Indeed, within minutes after the injection, the blastoderm exhibited a sudden movement of approximately 60μm towards the posterior end of the embryo (**Fig. 3a** & Supplementary Video 9) consistent with the theoretical prediction that the posterior part of the embryo is pulling on the anterior part. This confirms that the blastoderm-vitelline contact is indeed sustaining a substantial force.

**Figure 3:**
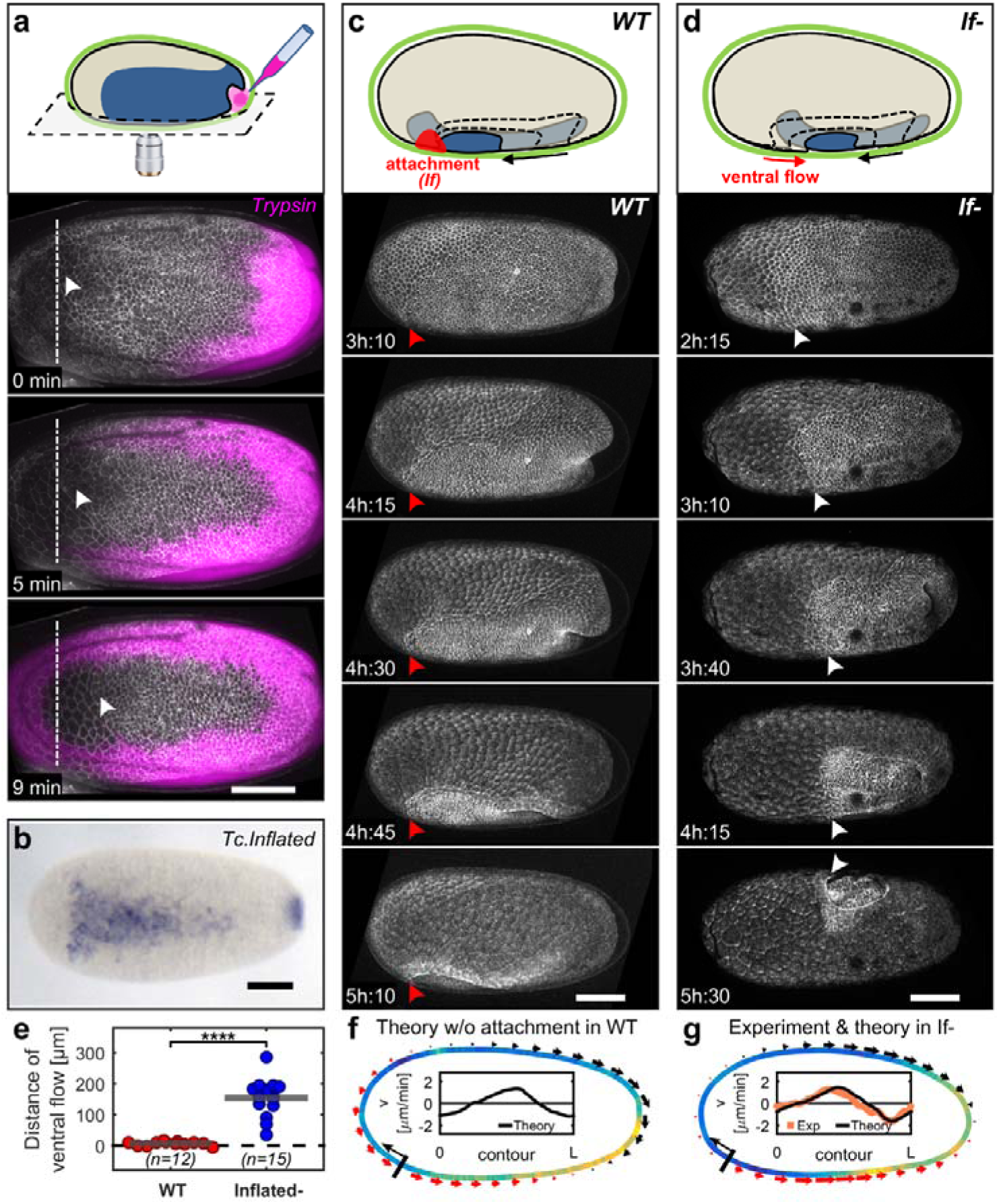
Release of blastoderm attachment causes ectopic ventral tissue flows. (**a**) Ventral view of a Tribolium embryo expressing LifeAct-eGFP after a solution of T rypsin containing a trace amount of rhodamine-dextran (magenta) was injected into the perivitelline space at the posterior pole (top panel schematic). The border between the serosa and embryo is marked by a dashed line and arrowheads trace the sudden movement of the tissue towards the posterior. (**b**) Ventral view of an embryo stained by *in situ* hybridization with anti-sense RNA for *Tc.inflated.* (**c**) Lateral view of a spinning disc confocal time-lapse of an embryo injected with LifeAct-eGFP mRNA. Red arrowheads mark the stationary area of the blastoderm. Top panel shows the process schematically marking the putative attachment zone in red and flow direction (black arrow). (**d**) Lateral view of a spinning disc confocal time-lapse of an embryo injected with mRNA encoding LifeAct-eGFP and dsRNA against *Tc.inflated* (If-). White arrows mark ectopic flow of the ventral border between serosa and embryo towards the posterior. Top panel shows the mutant phenotype schematically marking flows by red and black arrows. Scale bars in **a-d**: 100μm. (**e**) Quantification of the ectopic ventral flow towards the posterior in *Tc.inflated* knock-down embryos compared to wild type. Horizontal bars denote the mean of the distributions (pval=1.24^∗^10^-5^ Wilcoxon rank sum test). (**f**) Predictions of theory without attachment in wild type embryos. (**g**) Predictions of theory without attachment in *Tc.inflated*^-^ embryos. The flow profile predicted by a thin-film active fluid theory with no attachment matches the measured flow (inset and also compare to **f**).

Since non-specific digestion of proteins in the perivitelline space released the blastoderm attachment and TEM data showed a possible protein complex at the tip of the microvilli attached to the vitelline, it is conceivable that a specific molecular machinery rather than passive mechanical coupling and unspecific friction mediates the interaction. To investigate this, we searched for proteins that would be zygotically expressed specifically in the attachment zone, mediating some form of cell to extracellular matrix interaction. Possible candidate proteins that could fulfil such function are integrins, a class of transmembrane proteins that attach cells to the extracellular matrix^36^. Indeed, the expression pattern of the homolog of Drosophila alphaPS2-integrin *inflated* (*if*), (TC001667 *Tc.inflated*), coincides precisely with the attachment zone in the ventral blastoderm^37^ (**Fig. 3b**). We next tested whether *Tc.inflated* plays a role in attachment of the blastoderm to the vitelline envelope by downregulating the gene in the embryo using RNA interference^38^ (RNAi) and determining the effect on tissue flow by live imaging. The results showed that embryonic RNAi knockdown of *Tc.inflated* caused abnormal ventral tissue flow towards the posterior of the embryo during otherwise wild-type gastrulation (**Fig. 3c-e** & Supplementary Videos 10,11). Remarkably, this ectopic ventral flow matches the initial prediction of an active thin film fluid theory without the attachment point that we had initially applied to wild-type (**Fig. 3f**). In addition, when we predicted tissue flows from the myosin distribution imaged with light sheet in *Tc.inflated* RNAi embryos, the theoretical description without an attachment quantitatively matched our observations (**Fig. 3g** & Supplementary Video 12). Thus, in detached embryos, tissue-intrinsic forces alone are sufficient to explain the tissue flows. Moreover, the strength of the observed *Tc.inflated* knockdown phenotype varied and in extreme cases the embryos rotated inside the egg shell resulting in a random positioning of the serosa window and the embryo (**Fig. 3d** & Supplementary Videos 10,11). This observation further corroborates a complete loss of attachment to the vitelline. In conclusion, our data are consistent with the blastoderm being attached to the vitelline envelope by an alphaPS2-integrin, and that this attachment is necessary for normal gastrulation in Tribolium.

We next wondered if the integrin-mediated interaction of blastoderm and vitelline observed in Tribolium could be a general feature of insect gastrulation. Interestingly, the expression of alphaPS integrins in the fruit fly *Drosophila melanogaster* is restricted to specific regions of the blastoderm prior to gastrulation^39^. Among these, the alphaPS3 integrin *scab* (*scb)* is zygotically expressed in the posterior dorsal region of the blastoderm where the hindgut primordium invaginates during gastrulation^40^ (**Fig. 4a**). We set out to test if, similar to Tribolium, this integrin might mediate an interaction of the hindgut primordium with the vitelline. Indeed, a blastoderm vitelline proximity map (**Fig. 4b** & Supplementary Video 13) and a trypsin digestion experiment (Supplementary Video 14) are consistent with a force bearing contact between the hindgut primordium and the vitelline envelope during germ-band extension. In the light of the Tribolium results, we predicted that abolishing such an interaction would cause abnormal gastrulation movements, and we indeed observed that the germ-band dramatically twisted to the lateral side in *scb* mutants^40^ (**Fig. 4c,d &** Supplementary Videos 15,16). This result suggests that Scab plays a role in keeping the hindgut aligned with the dorsal midline of the embryo, possibly by mediating frictional forces between blastoderm and vitelline.

**Figure 4:**
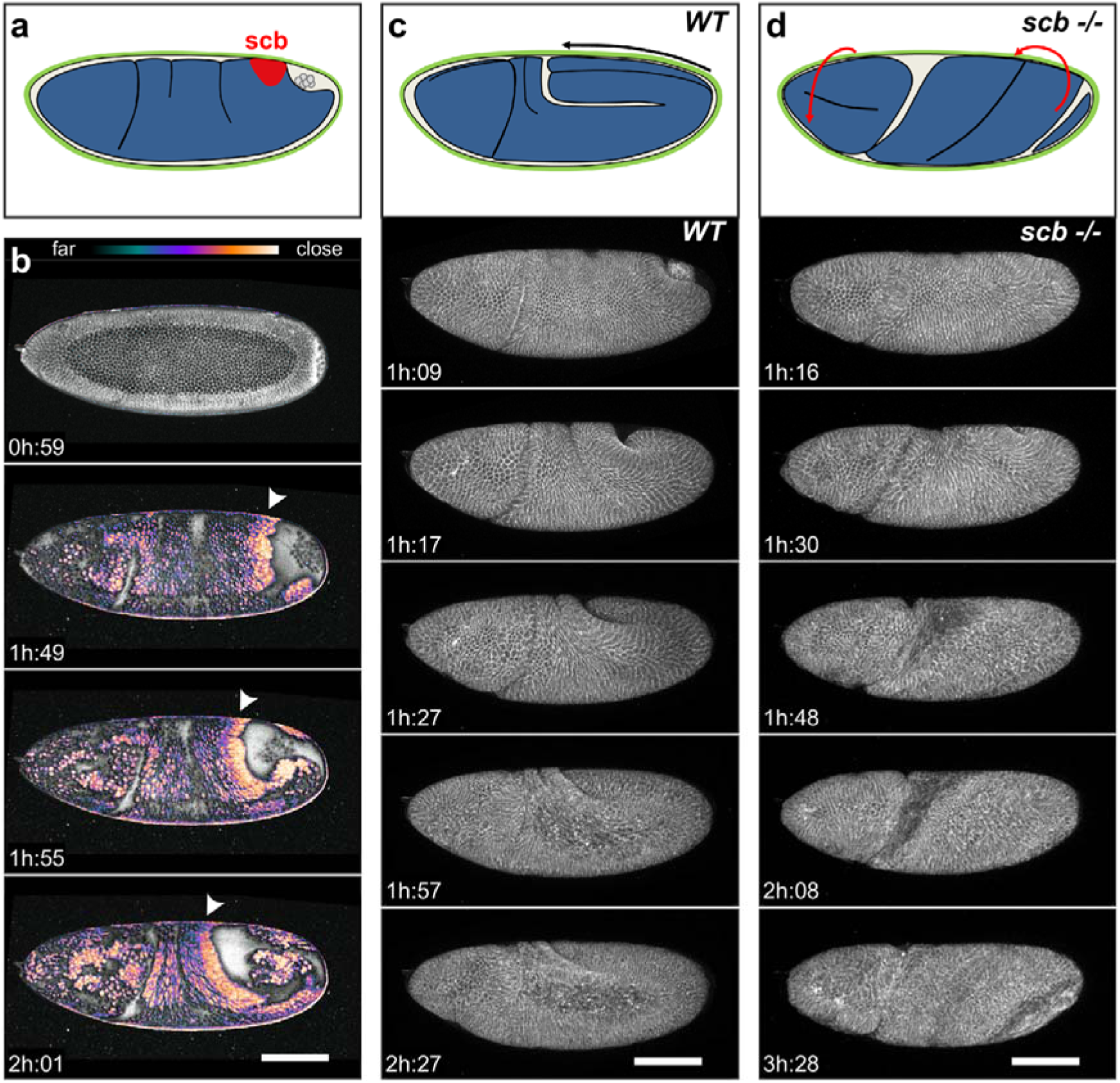
Integrin-mediated tissue adhesion is also necessary for normal germband extension in *Drosophila melanogaster.* (**a**) Schematic representation of Drosophila embryo (blastoderm in blue) at the beginning of germ band extension: the embryo is surrounded by the vitelline envelope (green), pole cells mark hindgut invagination, the approximate location of *scb* mRNA expression^40^ is highlighted in red. (**b**) A blastoderm vitelline proximity map (analogous to **Fig. 2g** in Tribolium) of Drosophila embryo expressing GAP43-mCherry. Arrowheads mark the anterior lip of hindgut invagination. (**c**) Lateral view of Drosophila germ band extension imaged using a confocal microscope in a strain expressing GAP43-mCherry. Top panel shows fully extended germ band schematically; arrow indicates the direction of the tissue movement. (**d**) Lateral view of scb^2^ homozygous mutant Drosophila embryos expressing GAP43-mCherry revealing germband twist to the lateral side while the head exhibits a counter twist. The mutant phenotype is schematically depicted in the top panel with directions of tissue rotations marked by red arrows. Scale bars in **b-d**: 100μm.

Taken together, the application of a theory relating myosin contractility to tissue flow in Tribolium gastrulation revealed a tissue-extrinsic force that was previously not considered relevant for tissue mechanics in insects. This force originates from a spatiotemporally regulated pattern of adhesion between the blastoderm and the surrounding vitelline envelope. The resulting anchoring is likely mediated by integrins and counteracts tissue-intrinsic contractile forces to generate asymmetric flows during gastrulation of Tribolium. The pattern of interaction between blastoderm and vitelline is different in Tribolium and Drosophila, giving rise to the intriguing possibility that the evolution of the regulation of integrin expression within the blastoderm is one of the mechanisms contributing towards diversification of early embryonic morphogenesis in insects^41^. In the future, it will be interesting to investigate the exact molecular mechanism and the spatio-temporal control of blastoderm adhesion to the vitelline as well as its impact on the evolution of early morphogenetic processes in insects and other organisms.

## Methods

### Stock keeping & transgenic lines

*Tribolium castaneum* (Herbst) stocks were cultured on organic whole wheat flour with dry yeast powder added^42^. Stocks were maintained at 32°C and 70% relative humidity. All mRNA injection experiments were conducted as described previously^18^ with *Vermillion white* mutants. A transgenic line constitutively expressing αTub:LifeAct-eGFP was used to visualize F-actin during gastrulation^43^.

To characterize the phenotype of *Drosophila melanogaster scb* mutants we combined by genetic crossing fly strains carrying (i) homozygous *scb* alleles (cn^1^ scb^2^ bw^1^ sp^1^/CyO; (Bloomington stock Nr. 3098) and cn^1^ P{PZ}scb^01288^/CyO; ry^506^ (Bloomington stock Nr. 11035)), (ii) GAP43 membrane marker (w^1118^; Gap43-mCherry/TM3Sb^44^) and (iii) a fluorescently labelled balancer on the second chromosome. Progeny of the resulting fly strains (w^1118^; scb^2^ or ^01288^/CyO-GFP; Gap43-mCherry/+) were used for imaging of gastrulation phenotypes and homozygous mutant *scb* embryos were identified by the lack of GFP expression later during development. Flies were kept under standard conditions on standard medium.

### Plasmid generation and RNA synthesis

To visualize myosin, we amplified the full-length regulatory light chain of non-muscle myosin II (*spaghetti squash*, TC030667) from Tribolium cDNA by PCR and cloned it into pCS2+ vector to generate a GFP fusion (pCS2+-Tc.sqh-eGFP). Capped mRNA was subsequently transcribed *in vitro* with the mMessage machine Kit pCS2 (Invitrogen) according to the manufacturer’s instructions. Similarly, a pCS2+-syn21::LifeAct-eGFP plasmid (syn21 sequence from^45^) was used to generate LifeAct-eGFP mRNA for co-injection with dsRNA.

For RNAi silencing of *Tc.inflated*, dsRNA against two different regions of the *Tc.inflated* gene (TC001667) were used: iBeetle RNAi (iB_03227) was acquired from Eupheria Biotech GmbH, Dresden, Germany. The second dsRNA was made by *in vitro* transcription of a PCR product generated from Tribolium cDNA with the Megascript T7 Transcription Kit (Invitrogen). The gene region suitable for RNA interference was identified using DEQOR^46^ and amplified with the following primers: T7-promoter-CCGGTGCCGTCGAATAAAG and T7-promoter-GAAGTGCGATGCGTTTGATTG.

### Embryo injections

For injection experiments, *Vermillion white* or LifeAct-eGFP embryos were collected for two hours and after one additional hour dechorionated for 2 x 45 seconds in 20% v/v bleach. Carefully rinsed dechorionated embryos were injected with an Eppendorf Femtojet setup on a brightfield upright microscope. dsRNAs against *Tc.inflated* (final concentration of 1mg/ml) were mixed with LifeAct-eGFP mRNA (final concentration of 500ng/ml) to monitor the efficiency of injection. Tc.sqh-eGFP mRNA was injected at 1-2 mg/ml alone or mixed with *Tc.inflated* dsRNA at final concentration of 1mg/ml. For subsequent confocal imaging experiments, embryos were submerged in halocarbon 900 oil (Sigma) to prevent desiccation. For light-sheet imaging experiments, embryos were kept in a humidified petri dish until embedding in agarose.

For dextran injection into the perivitelline space of Tribolium, a trace amount of 70k MW tetramethylrhodamine-dextran (Invitrogen) was dissolved in ddH_2_0 and injected into the perivitelline space at the posterior pole of LifeAct-eGFP embryos approximately 6 hours after egg collection.

For trypsin digestion experiments, injections were conducted on the stage of a spinning disc confocal microscope (see section live imaging). A solution of 1x Trypsin (Sigma) in PBS containing a trace amount of rhodamine-dextran was injected into the perivitelline space at the posterior pole of a LifeAct-eGFP embryo and imaged immediately. As a control, an equivalent experiment was carried out by injecting only a diluted rhodamine-dextran solution (see Supplementary Video 10).

For Drosophila, a solution of FITC-dextran (Sigma) or 10 x Trypsin containing a trace amount of FITC-dextran was injected into the perivitelline space of GAP43-mCherry expressing embryos before the onset of gastrulation and during germband extension, respectively, followed by live imaging on a confocal microscope.

### Live imaging with spinning disc confocal and light-sheet microscope

Live imaging was carried out either on an Olympus - IX83, inverted stand confocal microscope equipped with a Yokogawa CSU-W1 spinning disc scan head, or on Zeiss LSM 710 NLO point scanning confocal system and on a Zeiss Microscopy GmbH Lightsheet Z.1 light sheet microscope. The microscopes were equipped with environmental control and experiments were conducted at 25°C or 30°C. On the Olympus-IX83 confocal microscope, an Olympus UPlan SApochromat 20x 0.4 or Olympus UPlan SApochromat 10x 0.4 lens was used for whole embryo time-lapse imaging yielding a pixel resolution of 0.59μm or 1.18 μm, respectively. Image stacks of 40 to 60 slices and a z-spacing of 3 μm were acquired overnight at a time interval of 3 to 6 minutes. For high-resolution imaging of the actin protrusions, either an Olympus UPLSAPO 60x 1.3 SIL lens was used and image stacks were acquired at a time interval of 10 seconds (**Fig. 2e**) or an Olympus UAPON 150x 1.45 TIRFM lens was used to obtain image stacks of the apical cell surface at 5 seconds time intervals (**Fig. 2d**). On the Zeiss LSM 710 microscope a 25x multi-immersion Zeiss LD LCI Plan-Apochromat 25x 0.8 Oil/Glyc/Water DIC objective was used with varying zoom yielding pixel resolutions between of 0.55 μm or 0.67 μm. On the light sheet microscope, a Zeiss Plan Apochromat 20x 1.0 W DIC lens was used together with pre-zoom of 0.6 resulting in a pixel resolution of 0.381 μm. Three or five regularly separated views of the embryos were acquired as image stacks of 90 to 100 z-planes at 2-3 μm spacing. The frame rate of the multi-view recordings (the time interval between beginnings of acquisitions of the first view) ranged between 90 seconds and 5 minutes. Specimens were mounted for multi-view light sheet microscopy in 1% low-melting point agarose dissolved in 1x PBS medium together with fluorescent beads (Estapor, Millipore) as described previously^47^. Tribolium imaging was done using either light sheet or spinning disc confocal and Drosophila imaging was done on the point scanning or spinning disc confocal microscope.

The time stamps for all experiments are with respect to the last synchronous round of nuclear divisions of the blastoderm in the respective species (time = 0 minutes) except for the trypsin injection experiment where the time of injection sets 0 minutes.

### Image processing

The multi-view light sheet data (referred to as SPIM data) were registered and fused in Fiji^48^ as previously described^24,47^. A custom developed plugin for BigDataViewer^49^ allowed the repositioning of the data to enable extraction of an image stack between 60 and 80 slices centered around the sagittal midplane of the embryo. This sub-stack of fused multi-view dataset was saved as a TIF image stack for subsequent analysis. First, a MATLAB (2015a, MathWorks) script implementing an active open contour algorithm was used to trace the embryo circumference for each time point (see Supplementary Video 1). For each pixel of this outline, the intensity along the inwards-pointing normal direction to the contour was summed up to yield a planar summed intensity projection (SIP) along the sagittal circumference (see Supplementary Video 1). This video sequence of SIPs capturing the myosin dynamics along the sagittal circumference of the embryo was further processed in MATLAB: (i) At each location along the circumference of the contour the average of the myosin intensity of all slices forming the sagittal sub-stack of the multi-view data was calculated. (ii) However, since the initial intensity profile of myosin can suffer from inhomogeneity owing to uneven mRNA distribution after injection, we normalized the intensity of all time points by the intensity profile of the initial time point. (iii) To further correct for possible intensity variation arising from varying illumination over time, the intensity profiles were normalized by the average intensity of benchmark fluorescent beads in the vicinity of the sample. This processing pipeline yielded the myosin intensity kymograph shown in Figure 1c. The one-dimensional time series of the normalized myosin intensity is used to solve Eq
. 6 (see Supporting Materials and Methods) for each time point. For purpose of displaying, the image data were adjusted for contrast and brightness at every time point in the movies.

To obtain the tissue movements at every time point, we used MATLAB to determine the displacements of image areas of a width of 32 pixels along the direction of the circumference following a standard approach of particle image velocimetry (PIV). In short, the cross-correlation between the 32 pixel wide template image of one time point and a larger image region around the same location of the subsequent time point was calculated. Then, the location of the highest value of the resulting cross-correlation was taken as the magnitude of displacement of this image area. The flow velocity of each image area is then given by the magnitude of their displacement divided by the frame interval. To achieve sub-pixel accuracy, template and search area were interpolated onto a 10-fold finer pixel array before calculation of the cross-correlation. The resulting one-dimensional flow field of tissue movements is smoothed with an interpolating spline function to even out PIV detection errors, which mainly arise from insufficient brightness of the template image. To check the validity of the obtained results, the flow field was overlaid with the intensity kymograph, as shown in Figure 1c. Qualitatively the calculated flow fields followed intensity features in the kymograph over time.

Cylindrical projections of the LifeAct-eGFP SPIM data were produced for better overview of all tissue movements with a custom Python script. In short, after fusion of the SPIM data, the anterior-posterior (AP) axis of the embryo was defined manually. At every pixel along this axis, the maximum intensity along evenly spaced radial lines was determined yielding a flattened projection of the 3D data. The resolution along the AP axis equals that of the original SPIM data; the resolution in the orthogonal direction, however, depends on the distance at which the maximal intensity is detected, and therefore, is not defined.

To produce a “proximity map” of the distance between blastoderm tissue and vitelline envelope, the dextran channel was processed as follows: The signal was first inverted and the histogram was then adjusted to yield a uniform zero before gastrulation and a high signal during gastrulation. Then a colormap was applied and the result was overlaid with the grayscale LifeAct-eGFP signal. All processing was restricted to the inside of the embryo by an appropriate mask.

Measurement of the extent of ventral flow for Figure 3e was carried out manually in Fiji by examining time-lapse confocal recordings of LifeAct-eGFP embryos and *Vermillion white* embryos injected with *Tc.inflated* dsRNA/LifeAct-eGFP mRNA. The position of the actin bridge connecting the forming head lobes was chosen manually as a reference position for the attachment point. The distance of this position during early blastoderm stage and the position of the same group of cells right before the serosa window closure was plotted as a measure of ventral flow distance.

### Theoretical analysis

The details of all theoretical derivations as well as the fitting procedure are given in the Supporting Materials and Methods document.

### TEM Tomography

Un-dechorionated LifeAct-eGFP embryos of appropriate stage were high-pressure frozen (Leica EM ICE) and automatically freeze-substituted (Leica EM AFS2) in a morphology cocktail containing acetone, 1% Osmiumtetroxide and 0.1% Uranylacetate. At room temperature, samples were gradually infiltrated with LX112-resin (Ladd Research) and polymerised at 60 °C. For tomography, 300 nm sections were cut with an Ultramicrotome (Leica Ultracut), post-contrasted with Uranyl acetate and lead citrate, and labelled with 15 nm gold fiducials. Tomograms were acquired at a 300 kV transmission electron microscope (Tecnai F30, equipped with camera Gatan “Ultrascan”), and reconstructed with help of the IMOD software^50^ (http://bio3d.colorado.edu/imod/).

### RNA *in situ* hybridization

Staged *Vermillion white* embryos were dechorionated and fixed to visualize the expression pattern of *Tc.inflated* by RNA *in situ* hybridization with digoxigenin-UTP-labeled RNA probes following a standard protocol^51^. The protocol was modified by including a finer rehydration series (PBS:methanol 3:7, 5:5, 7:3), longer washing steps (30 minutes each extending the protocol to 3 days) and hybridization buffer as described previously^52^. The anti-sense probe was transcribed with a T7 RNA polymerase from a PCR template amplified from cDNA with the following primers: T3-promoter-ACCAACACACGCTACAACCA and T7-promoter-ACCCACAAAGGCACAGTTTC.

## Supplementary Information

See SI Guide.

## Acknowledgments

We thank M. van der Zee for sharing the transgenic LifeAct-eGFP line, Y. Hsieh for LifeAct-eGFP mRNA and H2A-mCherry mRNA, P. Mejstrik for RNA in situs, T. Pietzsch for implementation of custom BigDataViewer plugin, the MPI-CBG EM facility for TEM images, M. Burkon and the MPI-CBG LMF for technical assistance, M. Benton, K. Panfilio and P. Gross for helpful discussions, and the Tribolium research community for support. We thank C. Norden and E. Knust and C. Zechner for careful review of the manuscript. A. J. was a member of the IMPRS-CellDevoSys PhD program and supported by a fellowship by the DIGS-BB. S.M. was supported by an ELBE post-doctoral fellowship.

## Author contributions

S.M. designed the research, performed experiments, analyzed the data, and wrote the manuscript. A.J. produced reagents and performed experiments. A.M. developed the theory and analyzed the data. A.P. suggested the project and produced reagents. S.W.G. and P.T. conceived and oversaw the project, and wrote the manuscript.

## Supplementary video captions

**Supplementary Video 1.** *Dimensionality reduction for theory.* Overview of dimensionality reduction required for theoretical analysis of blastoderm tissue flow. (upper left) 3D rendering of fused multi-view SPIM recording of a Tribolium embryo injected with Tc.sqh-eGFP. Red rectangle marks the location of the sagittal section, blue line highlights the embryo contour. (upper right) 2D sagittal cross-section through the SPIM data. The outline of the embryo (blue) is traced with an active open contour for each time point. (lower panel) 1D projection of the myosin intensity along the circumference of the sagittal cross-section from the upper right panel (from 0 to L (contour length)). The origin 0 is annotated in the upper right panel.

**Supplementary Video 2.** *Comparison of experiment and theory without attachment.* Comparison of the experimentally determined tissue flow profile with the theoretically predicted profile obtained by a thin-film active fluid theory that considers the tissue to flow freely with respect to the outer vitelline envelope (i.e. the model with constant homogenous friction, without attachment).

**Supplementary Video 3.** *Comparison of experiment and theory with attachment.* Comparison of the experimentally determined tissue flow profile with the theoretically predicted profile obtained by a thin-film active fluid theory that considers a local attachment of the tissue to the outer vitelline envelope at an anchor point.

**Supplementary Video 4.** *Dynamics of apical protrusions in Tribolium blastoderm.* Time-lapse recording of the apical region of a wildtype Tribolium embryo constitutively expressing LifeAct-eGFP by spinning disc confocal microscopy depicts the dynamics of apical actin protrusions in a dorsal region of the embryo during tissue movement.

**Supplementary Video 5.** *Dynamics of blastoderm vitelline attachment in Tribolium.* Time-lapse recording by spinning disc confocal microscopy of the ventral region of a wildtype Tribolium embryo expressing LifeAct-eGFP at the onset of gastrulation visualizes the dynamics of the contact between apical actin protrusions and the vitelline envelope. (upper panel) Maximum intensity projection of the x-y planes covering 5 μm approximately 20 μm deep inside the embryo. The yellow box outlines the region from which the y-z planes are taken for the lower panel. (lower panel) Maximum intensity projection of y-z planes covering approximately 12 μm. The red line was traced by an active open contour method to highlight the location of the vitelline envelope. While the apical side of the blastoderm tissue is initially well separated from the vitelline, some cells broaden their contact area and their distance decreases significantly over the course of gastrulation (white arrows).

**Supplementary Video 6.** *Blastoderm vitelline proximity map in Tribolium.* A time-lapse recording of a LifeAct-eGFP Tribolium embryo injected with a solution of rhodamine-dextran into the perivitelline space before gastrulation visualizes the dynamic changes in proximity between blastoderm tissue and vitelline envelope. The ventral side of the embryos was imaged with spinning disc confocal microscopy and visualized as raw maximum intensity projection of dextran signal (upper panel) and as inverted “proximity map” (color) highlighting the areas of dextran exclusion and signifying blastoderm contact to the vitelline envelope (lower panel).

**Supplementary Video 7.** *Attachment rip*-*off during serosa window closure in Tribolium.* Ventral region of a Tribolium embryo expressing LifeAct-eGFP shown in a cylindrical projection from reconstructed multi view SPIM recording depicts the sudden snap-off of the pinned edges of the anterior border of the serosa window (characterized by cable-like accumulation of actin) at a late stage of gastrulation.

**Supplementary Video 8.** *Lateral view of attachment peel*-*off during serosa window closure in Tribolium.* A time-lapse recording by spinning disc confocal microscopy of the ventral region of a Tribolium embryo expressing LifeAct-eGFP at the later stage of serosa window closure visualizes how the blastoderm tissue is forcibly pulled away from the vitelline by the closing serosa window (arrow). The first 15 frames are repeated 3 times at the end of the movie.

**Supplementary Video 9.** *Attachment disruption by Trypsin digestion in Tribolium.* A time-lapse recording by spinning disc confocal microscopy of the ventral region of a Tribolium embryo expressing LifeAct-eGFP (gray) injected with a solution of trypsin spiked with rhodamine-dextran (magenta) is repeated 2 times in the upper panel. The trypsin/dextran solution was injected into the peri-vitelline space at the posterior during gastrulation and immediately imaged demonstrating the release of attachment force by proteolytic digestion. (lower panel) Similar experiment with the injection of a solution of only rhodamine-dextran as a control shows no retraction after injection.

**Supplementary Video 10.** *Release of attachment by Tc.nflated RNAi in Tribolium I.* Comparison of the blastoderm movements of a Tribolium embryo injected with LifeAct-eGFP mRNA and a Tribolium embryo injected with a mixture of dsRNA against *Tc.inflated* and LifeAct-eGFP mRNA imaged by spinning disc confocal microscopy. (upper panel) Lateral view of a wildtype Tribolium embryo. (lower panel) Lateral view of an inflated-knockdown embryo. Strong ectopic movement of the ventral tissue towards the posterior is noticeable concurrent with the onset of dorsal blastoderm movement. At the end of the time-lapse, the embryo turns inside the egg shell before the serosa window is fully closed and consequently the developing embryo becomes ectopically positioned on the dorsal side of the egg.

**Supplementary Video 11.** *Release of attachment by Tc.inflated RNAi in Tribolium II.* Overview of knock-down phenotype in several different embryos injected with dsRNA against *Tc.inflated* and mRNA encoding for LifeAct-eGFP recorded by spinning disc microscopy. (left) Embryos injected with *Tc.inflated* dsRNA obtained from iBeetle resource at 25°C. (right) Embryos injected with home-made *Tc.inflated* dsRNA at 30°C.

**Supplementary Video 12.** *Comparison of experiment and theory in Tc.inflated knockdown embryos.* Comparison of the tissue flow profile experimentally determined from multi-view light sheet microscopy of a Tribolium embryo injected with Tc.sqh-eGFP mRNA and dsRNA against *Tc.inflated* with the theoretically predicted profile obtained by a thin-film active fluid theory that does not consider any tissue attachment. The prediction of attachment-less theory matches the observed *Tc.inflated* phenotype quantitatively.

**Supplementary Video 13.** *Blastoderm vitelline proximity map in Drosophila.* A time-lapse of a lateral view of a Drosophila embryo expressing GAP43-mCherry (grey signal) injected with a solution of FITC-dextran into the perivitelline space before gastrulation. The embryo was imaged with spinning disc confocal microscopy and visualized as raw maximum intensity projection of dextran signal (upper panel) and as inverted “proximity map” (color) highlighting the areas of dextran exclusion. In particular, the anterior lip of the involuting hindgut primordium shows strong exclusion signifying local contact of the blastoderm to the vitelline envelope (lower panel).

**Supplementary Video 14.** *Trypsin injection into the perivitelline space in Drosophila.* Posterior dorsal region of a wildtype Drosophila embryo expressing GAP43-mCherry (gray) was injected with a solution of Trypsin spiked with FITC-dextran (magenta) into the perivitelline space at the posterior during hindgut involution and immediately imaged on spinning disc confocal microscope to visualize the release of tissue attachment by proteolytic digestion.

**Supplementary Video 15.** *Twisted gastrulation phenotype in Drosophila embryos mutant for scab I.* Comparison of the gastrulation in Drosophila embryos homozygous for *scb^2^* mutant allele and expressing GAP43-mCherry, with heterozygous *scb^2^*/+ siblings showing GAP43-mCherry signal as a control. Embryos were recorded by spinning disc confocal microscopy. Upper panel shows lateral view of control Drosophila embryos exhibiting normal progression of germ-band parallel to the dorsal midline. Lower panel shows lateral view of a *scb* mutant embryo exhibiting a phenotype where the germ band twists off the axis defined by the embryo dorsal midline.

**Supplementary Video 16.** *Twisted gastrulation phenotype in Drosophila embryos mutant for scab II.* Time-lapse recordings by spinning disc confocal microscopy of the gastrulation phenotype of several Drosophila embryos deficient for Scab protein that are also constitutively expressing GAP43-mCherry to visualize embryo anatomy (gray). Left panels are showing phenotypes of *scab^01288^/scab^01288^* and right panels are showing phenotypes of *scab^2^/scab^2^*.

## References

1. Gastrulation: From Cells to Embryo. (Cold Spring Harbor Laboratory Press, 2004).

2. Roux, W. Gesammelte Abhandlungen über Entwickelungsmechanik der Organismen: Bd. Entwicklungsmechanik des Embryo. 2, (Wilhelm Engelmann, 1895).

3. Gustafson, T. & Wolpert, L. Cellular Movement and Contact in Sea Urchin Morphogenesis. Biol. Rev. 42, 442–498 (1967).

4. Gilbert, S. F. Developmental Biology, Tenth Edition. (Sinauer Associates, Inc., 2013).

5. Turner, F. R. & Mahowald, A. P. Scanning electron microscopy of Drosophila embryogenesis. 1. The structure of the egg envelopes and the formation of the cellular blastoderm. Dev. Biol. 50, 95–108 (1976).

6. Heisenberg, C.-P. & Bellaïche, Y. Forces in Tissue Morphogenesis and Patterning. Cell 153, 948–962 (2013).

7. Heer, N. C. & Martin, A. C. Tension, contraction and tissue morphogenesis. Development 144, 4249–4260 (2017).

8. Leptin, M. Drosophila Gastrulation: From Pattern Formation to Morphogenesis. Annu. Rev. Cell Dev. Biol. 11, 189–212 (1995).

9. Zallen, J. A. & Wieschaus, E. Patterned gene expression directs bipolar planar polarity in Drosophila. Dev. Cell 6, 343–355 (2004).

10. Bertet, C., Sulak, L. & Lecuit, T. Myosin-dependent junction remodelling controls planar cell intercalation and axis elongation. Nature 429, 667–671 (2004).

11. Rauzi, M., Verant, P., Lecuit, T. & Lenne, P.-F. Nature and anisotropy of cortical forces orienting *Drosophila* tissue morphogenesis. Nat. Cell Biol. 10, 1401–1410 (2008).

12. Martin, A. C., Kaschube, M. & Wieschaus, E. F. Pulsed contractions of an actin–myosin network drive apical constriction. Nature 457, 495–499 (2009).

13. Butler, L. C. et al. Cell shape changes indicate a role for extrinsic tensile forces in Drosophila germ-band extension. Nat. Cell Biol. 11, 859–864 (2009).

14. Behrndt, M. et al. Forces Driving Epithelial Spreading in Zebrafish Gastrulation. Science 338, 257–260 (2012).

15. He, B., Doubrovinski, K., Polyakov, O. & Wieschaus, E. Apical constriction drives tissue-scale hydrodynamic flow to mediate cell elongation. Nature 508, 392–396 (2014).

16. Lye, C. M. et al. Mechanical Coupling between Endoderm Invagination and Axis Extension in Drosophila. PLoS Biol. 13, e1002292 (2015).

17. Rozbicki, E. et al. Myosin-II-mediated cell shape changes and cell intercalation contribute to primitive streak formation. Nat. Cell Biol. 17, 397–408 (2015).

18. Benton, M. A., Akam, M. & Pavlopoulos, A. Cell and tissue dynamics during Tribolium embryogenesis revealed by versatile fluorescence labeling approaches. Development 140, 3210–3220 (2013).

19. Handel, K., Grünfelder, C. G., Roth, S. & Sander, K. Tribolium embryogenesis: a SEM study of cell shapes and movements from blastoderm to serosal closure. Dev. Genes Evol. 210, 167–179 (2000).

20. Hannezo, E., Dong, B., Recho, P., Joanny, J.-F. & Hayashi, S. Cortical instability drives periodic supracellular actin pattern formation in epithelial tubes. Proc. Natl. Acad. Sci. U. S. A. 112, 8620–8625 (2015).

21. Dicko, M. et al. Geometry can provide long-range mechanical guidance for embryogenesis. PLoS Comput. Biol. 13, e1005443 (2017).

22. Streichan, S. J., Lefebvre, M. F., Noll, N., Wieschaus, E. F. & Shraiman, B. I. Global morphogenetic flow is accurately predicted by the spatial distribution of myosin motors. eLife (2018). doi:10.7554/eLife.27454

23. Huisken, J., Swoger, J., Del Bene, F., Wittbrodt, J. & Stelzer, E. H. K. Optical Sectioning Deep Inside Live Embryos by Selective Plane Illumination Microscopy. Science 305, 1007 (2004).

24. Preibisch, S., Saalfeld, S., Schindelin, J. & Tomancak, P. Software for bead-based registration of selective plane illumination microscopy data. Nat. Methods 7, 418–419 (2010).

25. Schmied, C., Stamataki, E. & Tomancak, P. Chapter 27 - Open-source solutions for SPIMage processing. in Methods in Cell Biology (eds. Waters, J. C. & Wittman, T.) 123, 505–529 (Academic Press, 2014).

26. Strobl, F. & Stelzer, E. H. K. Non-invasive long-term fluorescence live imaging of Tribolium castaneum embryos. Development dev.108795 (2014). doi:10.1242/dev.108795

27. Marchetti, M. C. et al. Hydrodynamics of soft active matter. Rev. Mod. Phys. 85, 1143–1189 (2013).

28. Prost, J., Jülicher, F. & Joanny, J.-F. Active gel physics. Nat. Phys. 11, 111–117 (2015).

29. Jülicher, F., Grill, S. W. & Salbreux, G. Hydrodynamic theory of active matter. Rep. Prog. Phys. 81, 076601 (2018).

30. Mayer, M., Depken, M., Bois, J. S., Jülicher, F. & Grill, S. W. Anisotropies in cortical tension reveal the physical basis of polarizing cortical flows. Nature 467, 617–621 (2010).

31. Furneaux, P. J. S. & Mackay, A. L. The composition, structure and formation of the chorion and the vitelline membrane of the insect egg-shell. in The insect integument 157–176 (Elsevier Amsterdam, 1976).

32. Grünfelder, C. G.-J. Vom frisch abgelegten Ei zum Blastoderm: Untersuchungen zur Feinstruktur der frühen Embryogenese des Reismehlkäfers Tribolium confusum, Duval (Coleoptera, Tenebrionidae). (Albert-Ludwigs-Universität Freiburg im Breisgau, 1997).

33. van Drongelen, R., Vazquez-Faci, T., Huijben, T. A. P. M., van der Zee, M. & Idema, T. Mechanics of epithelial tissue formation. J. Theor. Biol. 454, 182–189 (2018).

34. Jain, A. Molecular, Cellular and Mechanical basis of Epithelial Morphogenesis during Tribolium Embryogenesis. (Technical University Dresden, 2018).

35. Young, P. E., Richman, A. M., Ketchum, A. S. & Kiehart, D. P. Morphogenesis in Drosophila requires nonmuscle myosin heavy chain function. Genes Dev. 7, 29–41 (1993).

36. Bökel, C. & Brown, N. H. Integrine in Development: Moving on, Responding to, and Sticking to the Extracellular Matrix. Dev. Cell 3, 311–321 (2002).

37. Stappert, D., Frey, N., Levetzow, C. von & Roth, S. Genome wide identification of Tribolium dorsoventral patterning genes. Development dev.130641 (2016). doi:10.1242/dev.130641

38. Posnien, N. et al. RNAi in the Red Flour Beetle (Tribolium). Cold Spring Harb. Protoc. 2009, pdb.prot5256 (2009).

39. Sawala, A., Scarcia, M., Sutcliffe, C., Wilcockson, S. G. & Ashe, H. L. Peak BMP Responses in the Drosophila Embryo Are Dependent on the Activation of Integrin Signaling. Cell Rep. 12, 1584–1593 (2015).

40. Stark, K. A. et al. A novel alpha integrin subunit associates with betaPS and functions in tissue morphogenesis and movement during Drosophila development. Development 124, 4583–4594 (1997).

41. Kalinka, A. T. et al. Gene expression divergence recapitulates the developmental hourglass model. Nature 468, 811–814 (2010).

42. Brown, S. J. et al. The Red Flour Beetle, Tribolium castaneum (Coleoptera): A Model for Studies of Development and Pest Biology. Cold Spring Harb. Protoc. 2009, pdb.emo126 (2009).

43. van Drongelen, R., Vazquez-Faci, T., Huijben, T. A. P. M., van der Zee, M. & Idema, T. Mechanics of epithelial tissue formation. J. Theor. Biol. 454, 182–189 (2018).

44. Martin, A. C., Gelbart, M., Fernandez-Gonzalez, R., Kaschube, M. & Wieschaus, E. F. Integration of contractile forces during tissue invagination. J. Cell Biol. 188, 735–749 (2010).

45. Pfeiffer, B. D., Truman, J. W. & Rubin, G. M. Using translational enhancers to increase transgene expression in Drosophila. Proc. Natl. Acad. Sci. U. S. A. 109, 6626–6631 (2012).

46. Henschel, A., Buchholz, F. & Habermann, B. DEQOR: a web-based tool for the design and quality control of siRNAs. Nucleic Acids Res. 32, W113–W120 (2004).

47. Schmied, C., Steinbach, P., Pietzsch, T., Preibisch, S. & Tomancak, P. An automated workflow for parallel processing of large multiview SPIM recordings. Bioinformatics 32, 1112–1114 (2016).

48. Schindelin, J. et al. Fiji: an open-source platform for biological-image analysis. Nat. Methods 9, 676–682 (2012).

49. Pietzsch, T., Saalfeld, S., Preibisch, S. & Tomancak, P. BigDataViewer: visualization and processing for large image data sets. Nat. Methods 12, 481–483 (2015).

50. Kremer, J. R., Mastronarde, D. N. & McIntosh, J. R. Computer Visualization of Three-Dimensional Image Data Using IMOD. J. Struct. Biol. 116, 71–76 (1996).

51. Fonseca, R. N. da et al. Self-Regulatory Circuits in Dorsoventral Axis Formation of the Short-Germ Beetle Tribolium castaneum. Dev. Cell 14, 605–615 (2008).

52. Tomancak, P. et al. Systematic determination of patterns of gene expression during Drosophila embryogenesis. Genome Biol. 3, research0088.1 (2002).

